# Gene Regulatory Networks that support Multi-Fate Cellular Decisions

**DOI:** 10.64898/2026.07.13.738161

**Authors:** Harshavardhan BV, Sarah Adigwe, Mohit Kumar Jolly, Tomáš Gedeon

## Abstract

Cell fate decisions are driven by gene regulatory networks (GRNs). While the mutually inhibitory toggle switch effectively models binary fate decisions, fully connected inhibitory networks with more than two nodes fail to capture multi-fate decisions due to the low prevalence of “single high states”, where only a single master regulator is highly expressed. The goal of this study is to find network structures that support all single high states. We find that the only network that attains the highest possible prevalence of all single high states within the set of monotone Boolean (MB) models is completely disconnected. Since biological networks typically require connectivity, we investigate network structures that support equipotency, where all single high states have equal prevalence within MB models. Finally, we characterize the networks that support multistability between all single high states, finding that it is possible only in networks in which each node either has self-activations or is inhibited by every other network node. Our findings provide a theoretical framework for understanding the network design principles that can support simultaneous differentiation into multiple distinct cell types.

## 1. Introduction

Gene regulatory networks (GRNs) regulate the expression patterns during development to determine cell fates, transitioning from stem-like states to differentiated states [30]. Transcription factors (TFs) form the core of these networks by interacting with other transcription factors and regulating downstream genes. A specific set of these transcription factors, known as master regulators [6], regulates the differentiation towards a particular cell state, and these differentiated cells are characterized by exclusive expression of the master regulator corresponding to that particular state. The lack of experimental evidence for the regulatory links between the master regulators poses a challenge for a bottom-up experimental approach to identify network structures. Since the precise network interactions and processes that lead to these differentiated states are not known, we ask a question about what network structures support multiple steady states, where in each state, only one master regulator is expressed at a high level, and others are at a low level. We call such states *single high states* of a network. We are interested in discovering what network, or networks, of interactions between master regulators will maximize the prevalence of single high states, and what networks admit multistability between all single high states.

Previous studies considered pairs of master regulators forming the network motif of a toggle switch, where the mutual inhibition between two TFs allows for mutually exclusive expression of both single high states [13, 27]. In our previous study [4], we extended this architecture and considered fully connected mutually inhibitory networks with more than two nodes. We hoped that such a network would support a multi-fate decision by supporting a high prevalence of all single high states. However, we observed by several complementary methods that such networks do not give rise to a predominance of single high states. This leaves an open question whether any other network structure would allow for the predominance of single high states.

To address this question, we first define a *prevalence* of a particular single high state as the number of monotone Boolean models that support such a state, normalized by all monotone Boolean models that are compatible with the network structure. Regulatory network can be represented as a directed graph *N* = *(V, E)* with vertices *V* and directed edges *E*, and, in addition, with a sign *δ (e)* ∈ {±1} associated to each edge in *E*. The sign determines whether the directed interaction is activating (*δ (e)* = 1) or repressing (*δ (e)* = −1). Since the monotone Boolean models that are compatible with the network structure depend on the number of nodes and the number of edges of the network, but not the signs of the edges, the proper comparison can be made between networks with the same directed edge structure, but that differ in the sign structure.

We therefore ask the question about the prevalence of single high states in the context of fixing the directed graph structure of the network, i.e., the nodes *n*, edges *E*, and then ask what sign structure *δ* on the set of edges supports multiple single high states with the highest possible prevalence. We further investigate two properties: equipotency and multistability. The first asks which sign structure supports all single high states with *the same number* of monotone Boolean models, where the models that support different states may not be the same. We say that the regulatory network is *equipotent* if it has this property. If we have no knowledge of what monotone Boolean model is the model for the network, an equipotent network would produce any single high state with equal probability under repeated sampling of compatible monotone Boolean models. The second question asks which sign structure of the network has at least one monotone Boolean model that supports *multistability* between all single high states. Once we find such structures, we can then attempt to rank them based on the number of such models.

The paper is organized as follows. After preliminaries in Section 2, Section 3 presents the main tool that allows us to study network structures that support particular sets of equilibria. We extend the characterization of monotone Boolean models for networks with positive edges supporting a particular equilibrium, presented in [2], to include all networks with both positive and negative edges. In Section 4 we describe for any network structure and any Boolean state *b* of the network the set of monotone Boolean models (MBM) that are compatible with that network structure and for which *b* is a fixed point. We show that the only network structure with *n* nodes that is able to support all *n* single high states with the highest possible prevalence is a completely disconnected network with positive self-regulation on each node.

Since networks without mutual communication cannot serve as decision-making circuits, in Section 5 we look at equipotent networks. We provide a characterization of such networks within a class of networks with all-to-all edges with no self-loops in Theorem 5.5 and then apply our results for classes of networks with *n* = 3, 4, 5 nodes. We find that the highest prevalence among the equipotent networks occurs for *n* = 3 when each node receives negative inputs from both other nodes, for *n* = 4 each node either receives one negative inputs or negative inputs from all three other nodes, and for *n* = 5 when each node receives one negative input. When we expand the study to look at networks with all-to-all coupling with self-regulation, the types of networks that maximize prevalence remain largely the same. However, the relationship between *n* and the number of negative inputs that support maximal equipotency for *n >* 5 remains an open problem.

Finally, in Section 6 we focus on network structures that support multistability between all *n* single high states. Remarkably, in Theorem 6.1 we find that the only networks that support such multistability must, for each node, either have positive self-regulation or the node receives negative input from all other nodes. We then ask which structures among these have the highest number of models that support multistability. For *n* = 3, 4, these are networks that have both of these features, positive self-regulation and negative input from all other nodes. However, for *n* = 5, this pattern changes, and the highest number of models is supported by a network with positive self-regulation on each node, but only one negative input from four other nodes. Full understanding of this pattern likely hinges on the detailed structure of the lattice of monotone Boolean functions and remains an open problem. We conclude with the discussion section.

## 2. Preliminaries

### Definition 2.1.

A *regulatory network N* = (*G, δ*) consists of

- a directed graph *G* = (*V, E*) with nodes *V* with |*V* | = *n* and directed edges *E*;
- *edge sign function δ* : *E* → {− 1, 1}.

We denote an edge from node *i* to node *j* without indicating its sign by (*i, j*) or *i* ⊸ *j*. The edge *i* ⊸ *j* is *activating* if *δ*(*i, j*) = 1 and *repressing* if *δ*(*i, j*) = − 1. Graphically, an activating edge of a regulatory network edge is denoted by *i* → *j* and a repressing edge by *i* ⊣ *j*. The *sources* and *targets* of a node *i* are given by

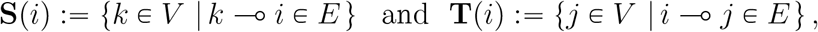

respectively.

Given a network *N*, we define a feasible set of models that represent the dynamics of the network. Because we are interested in the prevalence of certain dynamics within a class of models, we chose a class of models that is finite and thus enumerable. To do this, we start by assigning a Boolean state *b*_*i*_ ∈ B where B ≔ {0 < 1} has a state 0, representing the OFF state and a state 1, representing the ON state. Then the state of the network is a Boolean vector *b* ∈ B^*n*^ with *b* = *(b*_1_, …, *b*_*n*_) . The set (B^*n*^,⪯) is a partially ordered set under the natural order on vectors induced by the order 0 ⪯ 1 on B. We will use the notation ↑ (*b)* ≔ {*d* ∈ B^*n*^ |*b* ⪯ *d}* and ↓ (*b)* ≔ {*d* ∈ B^*n*^ | *d* ⪯ *b}* to denote *up* and *down* set of *b*.

The class of models that we consider is a class of *monotone Boolean models (MBM)* that we describe next.

A Boolean function *f*_*i*_ : B^*s*^ → B is <u>increasing with respect to input</u> *j* if for any input *b* = (*b*_1_, …, *b*_*s*_) ∈ B^*s*^

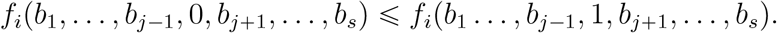

It is <u>strictly increasing with respect to</u> *j* if there is at least one input *b* ∈ B^*s*^ where the inequality is strict.

A Boolean function *f*_*i*_ : B^*s*^ → B is <u>decreasing with respect to input</u> *j* if for any input *b* = (*b*_1_, …, *b*_*s*_) ∈ B^*s*^

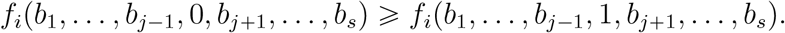

It is <u>strictly decreasing with respect to</u> *j* if there is at least one input *b* ∈ B^*s*^ where the inequality is strict.

### Definition 2.2.

A Boolean function *f*_*i*_ : B^*s*^ → B is <u>monotone</u> if it is increasing or decreasing with respect to each input *j =* 1, …, *s*. In this case, *f*_*i*_ is called a <u>monotone Boolean function</u>.

### Definition 2.3.

A Boolean model *f* : B^*n*^ → B^*n*^, *f* = *(f*_1_, …, *f*_*n*_), is <u>monotone</u> if for every node, *i* the function *f*_*i*_ is monotone.

A monotone Boolean model *f* : B^*n*^ → B^*n*^, *f* = *(f*_1_, …, *f*_*n*_), is <u>positive</u> (increasing) if for every *i*, the monotone Boolean function *f*_*i*_ is increasing with respect to all of its inputs.

### Definition 2.4.

A Boolean function *f*_*i*_ : B^*s*^ → B is redundant on input *j* if *f*_*i*_(*b*_1_, *b*_2_, ⃛, *b*_*j*−1_, 0, *b*_*j*+1_,⃛, *b*_*s*_) = *f*_*i*_(*b*_1_, *b*_2_, ⃛, *b*_*j*−1_, 1, *b*_*j*+1_, ⃛, *b*_*s*_) for all *b*_*k*_ ∈ B, *k* ≠ *j*. A Boolean function with no redundant inputs is called an *essential* Boolean function.

### 2.1. Network compatibility of Monotone Boolean models

Regulatory network *N* = *(G, δ)* consists of directed graph *G* = *(V, E)* and a sign function *δ* : *E* → *{*−1, 1}. For MBM *f* = *(f*_1_, …, *f*_*n*_), the directed graph structure determines the number of inputs of each monotone Boolean function *f*_*i*_ : B^|**S**(*i*)|^ → B. The sign structure *δ* determines the monotonicity type of *f*_*i*_, which we describe more formally here.

#### Definition 2.5.

A monotone Boolean model (MBM) *f* : B^*n*^ → B^*n*^, *f* =(*f*_1_, …, *f*_*n*_) is *compatible* with network *N* = (*G, δ*), *G* = (*V, E*) if |*V* | = *n* and for each (*i, j*), *f*_*j*_ is increasing (decreasing) with respect to *b*_*i*_ if, and only, if *δ*(*i, j*) = 1, (*δ*(*i, j*) = −1).

#### Definition 2.6.

There are two *elementary bijections* from B to B, identity *Id* and ⌐ *Id* which maps 0 → 1 and 1 → 0. For each vertex *i* ∈ *V*, let

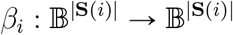

be a collection 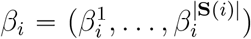 where each 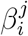 is an elementary bijection corresponding to the edge *j* ⊸ *i*.

The function *β*_*i*_ is determined by the sign function *δ*

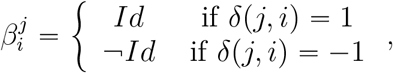

where *Id* is the identity function on B. The map 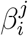 inverts Boolean input if the edge *j* ⊣ *i* is negative and preserves the Boolean input if the edge *j* → *i* is positive. We call *β* = (*β*_1_, …, *β*_*n*_), *β* : B^|*E*|^ → B^|*E*|^ a *change of sign* function.

Any sign function *δ* defines a unique Boolean sign function *β* and any Boolean sign functions *β* uniquely defines a sign function *δ*.

Clearly, the function *β*_*i*_ is determined by the set of negative input edges to node *i*. Let **Z**_*i*_ ⊂ **S**(*i*) be the set of nodes connecting by a negative edge to node *i*

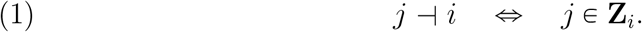

We use the shorthand 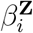 for the corresponding change of sign function.

### 2.2. Positive networks

Among the regulatory networks with the same *G* a *positive network* where all edges are positive (*δ(i, j)* = 1 for all (*i, j)*) has special properties. We start with an important definition.

#### Definition 2.7.

Consider the set *MBF* (*s*), a set of positive monotone Boolean functions *f* : B^*s*^ → B. For any input *b* ∈ B^*s*^ let

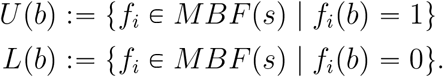

We denote a vector of zeroes in B^*n*^ by **0**, a vector of ones in B^*n*^ by **1**, and a *j*−th unit vector in B^*n*^ by *e*^*j*^, respectively. For any subset **Z** ⊂ {1, …, *n}* we denote by *e*^**Z**^ ∈ B^*n*^ the vector with 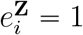 if, and only if, *i* ∈ **Z**. When it will be necessary to emphasize the size of these vectors, we will add a superscript, i.e., **1**^*s*^ ∈ B^*s*^.

We summarize some results from ([2]) about positive networks.

#### Theorem 2.8

(Theorem 2.10, [2]). *Consider a positive network N and an arbitrary Boolean state b* = *(b*_1_, …, *b*_*n*_) ∈ B^*n*^ *of the network N* .

*Then b is a steady state of any monotone Boolean model f* = *(f*_1_, *f*_2_, …, *f*_*n*_*) where*

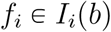

*where*

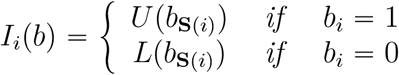

*where b*_**S**(*i*)_ *are the values of the input nodes of i evaluated at the state b*.

#### Theorem 2.9

(Theorem 2.16, [2]). *Consider a positive network N and q Boolean states b*^1^, …, *b*^*q*^ ∈ B^*n*^. *Then the multistability between these steady states is supported by any positive monotone Boolean model f* = (*f*_1_, *f*_2_, …, *f*_*n*_) *where*

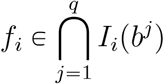

*where*

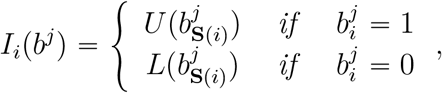

*where* 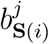 *are the values of the input nodes of i evaluated at the state b*^*j*^, *respectively*.

#### Theorem 2.10

(Theorem 2.12, [2]). *Consider a positive network N with n nodes*.

*Then*

- *The set of all MB models compatible with N is*

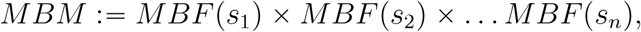

*where s*_*j*_ = |**S(***j)*|.
- *Every state in* B^*n*^ *is an equilibrium for some monotone Boolean model. Among all the states, the states* **0** ∈ B^*n*^ *and* **1** ∈ B^*n*^ *are most prevalent i*.*e. supported by most MBMs*.
- *The sets of MBMs that supports* **0** ∈ B^*n*^ *has the form*

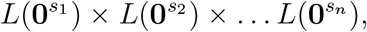

*which has the size*

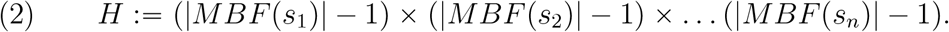
- *The sets of MBMs that supports* **1** ∈ B^*n*^ *has the form*

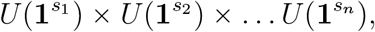

*which also has the size H*.

We have the following refinement of (2) that describes the sizes of all sets *U* (·) and *L*(·). For *b* ∈ B^*n*^ let |*b*| denote the number of 1s in Boolean vector *b*.

#### Lemma 2.11.

*(1)* |*U(b)*| = |*L(*⌐*b) for any b* ∈ B^*s*^

*(2) The size* |*U(b)*| *and* |*L(b)*| *depends only on* |*b*|. *In particular, there exists a set of integers*

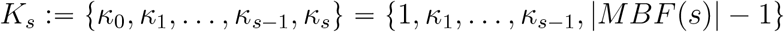

*with κ*_0_ < *κ*_1_ < …,*< κ*_*s*_ *such that for any b* ∈ B^*s*^

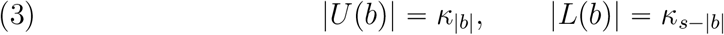

*Proof*. Statement (1) is Lemma 3.17 in [2] and (2) is Theorem 3.18 [2]. □

The set of numbers *K*_*s*_ is challenging to compute for arbitrary *s*. In particular, the size of the set of monotone Boolean functions |*MBF* (*s*)| is only known for *s* ⩽ 9 ([19]). Since *κ*_*s*_ ∈ *K*_*s*_ satisfies *κ*_*s*_ = |*MBF* (*s*)| − 1 computing the entire set *K*_*s*_ is at least as hard as computing |*MBF* (*s*)|.

## 3. Fixed points for general networks

We now describe our main theoretical result, where for any network *N*= *(G, δ)* and any *b* ∈ B^*n*^, we characterize the set of models that support *b* as a fixed point. We first observe that any monotone Boolean function *h*_*i*_ : B^|**S**(*j*)|^ → B which is monotonically decreasing with respect to its negative inputs and monotonically increasing with respect to its positive inputs can be written as a composition of a <u>positive</u> monotone Boolean function *f*_*i*_ : B^|**S**(*j*)| →^ B and the sign change function *β*_*i*_

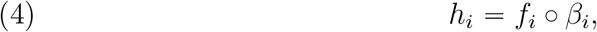

introduced in Definition 2.6. We have the following adaptation of Theorem 2.8 to general networks.

### Theorem 3.1.

*Consider an arbitrary network N* = *(G, δ) and an arbitrary Boolean state b* = *(b*_1_, …, *b*_*n*_ ) ∈ B^*n*^ *of the network N* . *Let β* = *(β*_1_, …, *β*_*n*_*) be the sign change induced by δ*.

*Then b is a steady state of any monotone Boolean model h* = (*h*_1_, *h*_2_, …, *h*_*n*_) *with h* = *f* ° *β, where*

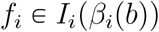

*where*

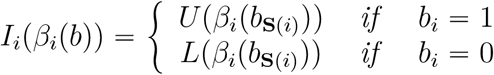

*where b*_**S**(*i*)_ *are the values of the input nodes of i evaluated at the state b*.

### Definition 3.2.

Fix network *N* =(*G, δ*) and associated function *β*.

The set of monotone Boolean models *supporting state e*^*i*^ ∈ B^*n*^ is the set

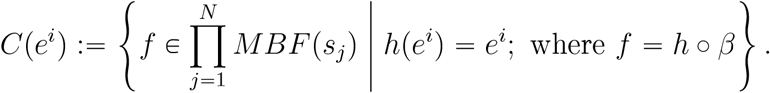

The regulatory network *N* is *equipotent* if

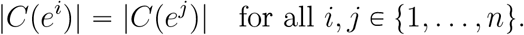

We have the following Corollary of Theorem 3.1 that characterizes the sets *C*(*e*^*i*^).

### Corollary 3.3.

*Let e*^*j*^ ∈ B^*n*^ *be a single high state. Then the set of MBM supporting e*^*j*^ *is*

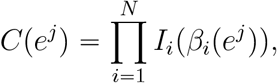

*where*

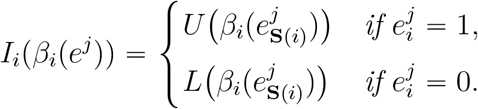

### Definition 3.4.

We say that a network *N* = *(G, δ)* supports *full multistability*, if there exists a MBM consistent with the network *N* for which all the single high states *e*^1^, …, *e*^*n*^ are steady-states of the network.

We have the following characterization of full multistability, which is a direct consequence of Theorem 2.9 for positive networks. This version incorporates the change of signs function *β*.

### Theorem 3.5.

*Consider network N* = *(G, δ). Then MBM h* = *(h*_1_, …, *h*_*n*_*) supports full multistability, if h* = *f ° β, where β is the sign change function associated to network sign structure δ and*

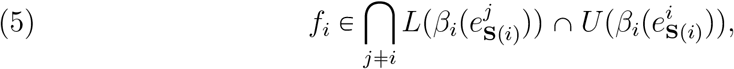

*where* 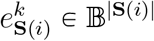 *is the binary vector of the values of the source nodes j* ∈ **S**(*i*), *evaluated at the steady state at e*^*k*^.

## 4. Maximum prevalence of single high states

We start with the following Corollary of Theorem 2.10 and (4).

### Corollary 4.1.

*Consider an arbitrary network N with n nodes and state space* B^*n*^. *Assume that the number of inputs to vertex i* ∈ *V is s*_*i*_ = |**S(***i)*|. *The highest possible number of monotone Boolean models that support an equilibrium e* ∈ B^*n*^ *is (see (2))*

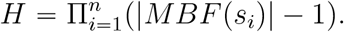

In view of this Corollary, we want to address the following question. We fix a particular state *e*^*i*^ ∈ B^*n*^ and ask which network among all networks with *n* nodes has the property that the size of the set of MB model that supports *e*^*i*^ as an equilibrium achieves the upper bound *H* in Corollary 4.1.

### Definition 4.2.

We say that network *N* = (*G, δ*) is a *k-team network* if there is a partition of the vertices *V* = *V*_1_ ∪ … ∪ *V*_*k*_ into *k* disjoint sets and

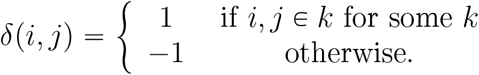

### Theorem 4.3.

*Consider state e*^*i*^ ∈ B^*n*^. *Then the only network with n nodes that supports e*^*i*^ *with H monotone Boolean models is a 2-team network where teams A, B are*

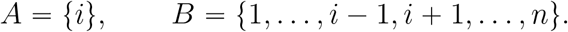

See Figure 1a for 2 node, Figure 1b for 3 node and Figure 1c the 4 node networks supporting *e*^1^.

*Proof*. By Theorem 3.1 the state *e*^*i*^ is supported by all models *f* = *(f*_1_, …, *f*_*n*_) such that

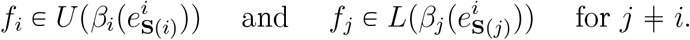

By Theorem 2.10 the only sets *U* (·) and *L*(·) with |*U* (·)| = |*L*(·)| = |*MBF* (·)|−1 are sets *U* (**1**) and *L*(**0**). Therefore, if *e*^*i*^ is supported by *H* MBMs then

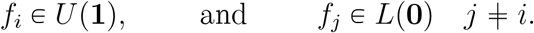

It follows that

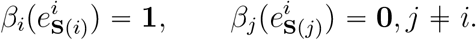

It follows from the condition 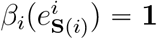 that

1. *i* → *i* (if it exists) is positive;
2. *j* ⊣ *i, j* ≠ *i* (if they exist) must be all negative.

On the other hand, condition 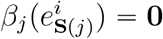 implies that

1. *i* ⊣ *j* (if it exists) must be negative;
2. *k* → *j, k* ≠ *i* (if they exist) must be all positive.

Since this holds for any *j* ≠ *i*, it must be that all existing edges from *A* = *{v*_1_} to any vertex *j* ∈ *B* ≔ *V\A* must be negative, edges from *B* to *A* must be negative, and edges between pairs of nodes in *B* are positive.

This proves the Theorem. □

**Figure 1.**
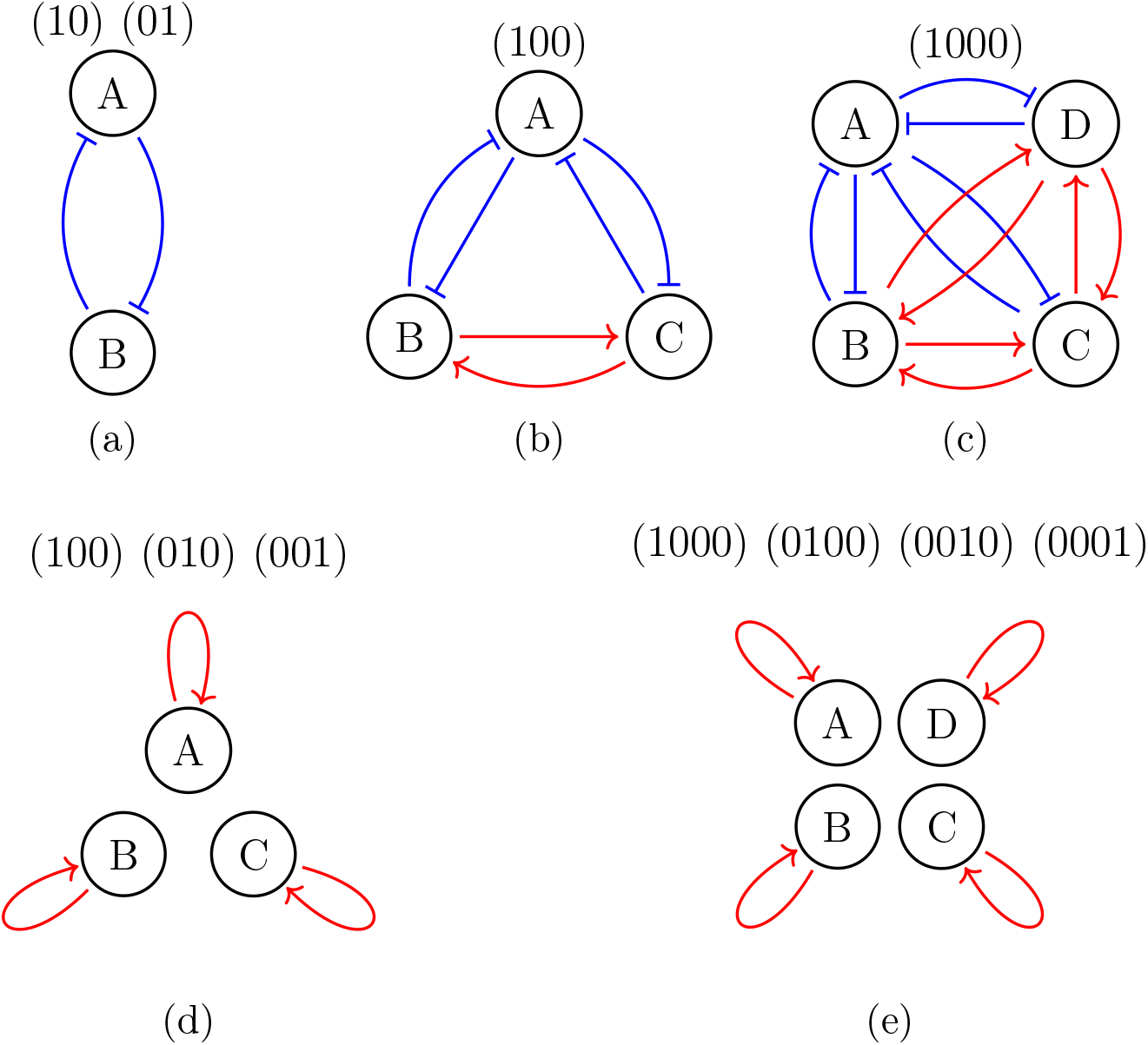
Network structure that support the highest prevalence (*H*) of single high states. (a)-(c) networks that support the state *e*^1^ with *H* MBMs: a two node network (a), a three node network (b), and a four node network (c). (d)-(e) networks that simultaneously support the state *e*^1^, …, *e*^*n*^ with *H* MBMs: a three node network (d), and a four node network (e). The states supported by *H* MBMs are indicated for each network.

### Corollary 4.4.

*The only n-node network N with n >* 2 *that supports every e*^1^, *e*^2^, …, *e*^*n*^ *with H monotone Boolean models is a completely disconnected network of n nodes, each of which has a positive self-edge, see Figure 1d for n* = 3 *and Figure 1e for n* = 4.

*Proof*. For *n* = 2 the toggle switch (see Figure 1a) is a network with teams *A* = {1} and *B* = {2} which supports *e*^1^ and *e*^2^ with the highest possible number *H* = 2 * 2 = 4 of MBMs.

.For *n* > 2, by Theorem 4.3, the network that supports *e*^*i*^ is the two team network *A*_*i*_ = {*i*} and *B*_*i*_ = *V*\*A*.

We now apply this observation to the collection of states *e*^1^, …, *e*^*n*^. Take any pair of nodes *i* ≠ *j*. Then *i* ∈ *A*_*i*_, *j* ∈ *B*_*i*_ which implies that if there is any edge between them, it must be negative. On the other hand, since *n* ⩾ 3, there is an index *k* ≠ *i, k* ≠ *j* and both *i, j* ∈ *B*_*k*_. This implies that if there is any edge between them, it must be positive. Since these conclusions are contradictory, we conclude that there is no edge in the network between any pair of nodes *i* ≠ *j*. Therefore, the only network that supports all single high equilibria *e*^1^, *e*^2^, …, *e*^*n*^ is a completely disconnected network of *n* nodes, each of which has a positive self-edge. □

### Remark 4.5.

Note that the domain of attraction of *e*^*i*^ in a fully disconnected network consists only of the state *e*^*i*^ itself.

## 5. Equal prevalence of single high states

Completely disconnected networks which support all the single high states with the highest possible number of models by Corollary 4.4 are not biologically meaningful. Therefore, we now relax this condition and ask which networks can support all *e*^1^, …, *e*^*n*^ by the same number of monotone Boolean models. This property has been defined in Definition 3.2 as equipotency. We do not require that this number be the highest possible number *H*.

We start by noting that for each component *f*_*j*_ of an equipotent MBM *f* = (*f*_1_, …, *f*_*n*_) has to satisfy constraints of Corollary 3.3 for all *e*^1^, …, *e*^*n*^. We assemble these constraints into a table

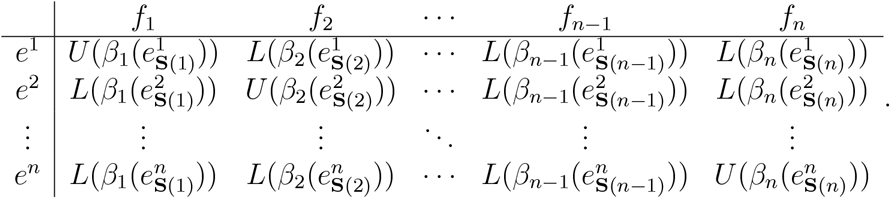

In this table, the entry in the *i*–th row and *j*–th column shows the set of functions *f*_*j*_ that support the equilibrium *e*^*i*^. Equipotency requires that the product of the sizes of the sets of models in each row be the same. In particular, consider a matrix of integer sizes

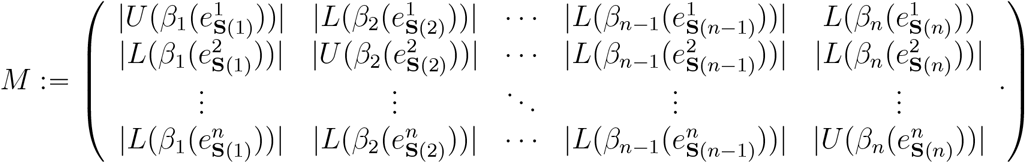

We have the following characterization of equipotency.

### Theorem 5.1.

*Consider network N* = *N (G, δ*) *and the function β corresponding to δ. Then the model f is equipotent for the network N* = (*G, δ*) *if the product of numbers in each row of M* = [*m*_*ij*_] *is the same i*.*e*.

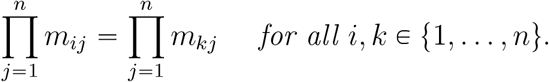

### 5.1. Networks with all-to-all coupling without self-edges

We now examine equipotency for general networks with all-to-all connections and no self-edges. We apply these results to networks with *n* = 3, 4, 5 nodes. We then extend these results to networks with self-edges. We show that the equipotency problem for such networks can be reduced to equipotency for networks with no self-edges.

Consider an *n* node network with all-to-all connections without self-loops. We ask which choice of edge signs *δ*, or equivalently, which choice of function *β* = (*β*_1_, …, *β*_*n*_) produces equipotent networks. We then select the maximum possible equipotent number achievable across all sign distributions *δ*.

While we cannot address this question in full generality, we can make substantial progress in simplifying the matrix *M* and find complete answers for *n* = 3, *n* = 4, and *n* = 5.

#### 5.1.1. General considerations

Consider networks *N* with directed graph *G* with *n* nodes and all directed edges except the self-edges.

We fix an order of vertices *V* = {1, …, *n*} . In the sequel, it will be useful to reorder the input set for each vertex *j*, **S(***j*) = {1, …, *j* − 1, *j +* 1, …, *n* } to be {1, …, *n* − 1 } in the order induced by the order on *V* .

Recall from preliminaries that **0** ^*n*−1^ denotes the zero vector and *e*^*k,n*−1^ denotes the *k*–th standard basis vector in B^*n*−1^. Then we have the following specification of the entries in the matrix *M*

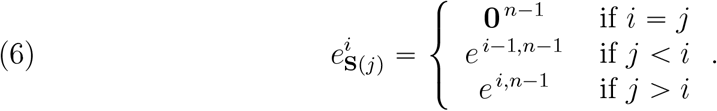

It follows immediately that for the positive network where *β*_*i*_ is the identity for all *i*, the matrix *M* for the network *N* takes the form

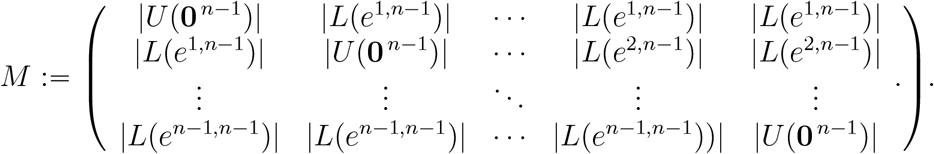

Let *B* = [*b*_*ij*_] be the matrix of inputs in the matrix *M* for the positive network:

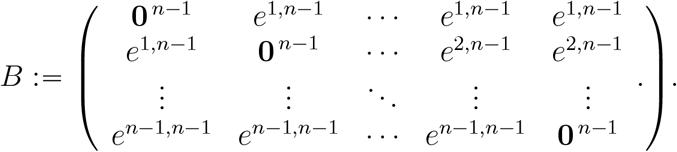

We discuss the effect of a general function *β* on the set of inputs by comparing the transformed set of inputs with the set of inputs for the positive network summarized in matrix *B*.

##### Theorem 5.2.

*Consider an n-node signed network with no self-edges. For each column j* ∈ {1, …, *n* }, *choose a subset* **Z**_*j*_ ⊆ **S (***j*) = {1, …, *n*−1 }, *of inputs into node j that correspond to negative edges*.

*Define the indicator vector of the set* **Z**_*j*_

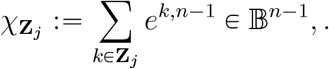

*Then, for every i, j* ∈ {1, …, *n*}

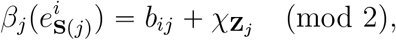

*where b*_*ij*_ *are elements of matrix B and the sum is modulo* 2.

*Proof*. Since each repressing edge replaces the corresponding input coordinate *x*_*l*_ by 1 − *x*_*l*_, the signed input configuration is obtained from the positive input vector by toggling selected coordinates. This is equivalent to the addition of the indicator vector (mod 2). □

The corresponding constraint matrix *M* has the form

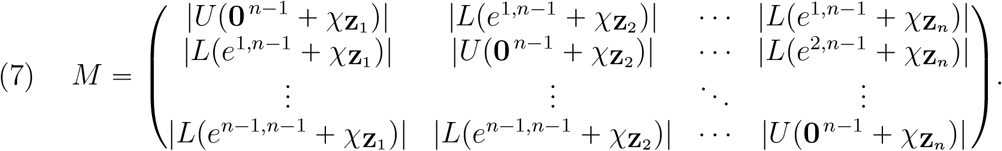

We now discuss how the arguments of the entries in matrix *M* depend on the choice of **Z**_*i*_, *i* = 1, …, *n*.

##### Lemma 5.3.

*Consider* |**Z**_*i*_| = *m. Then*

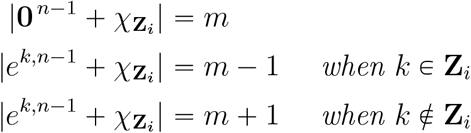

*Proof*. The first statement is immediate. On the other hand, for *i* ≠ *j* the entry *b*_*ij*_ = *e*^*k,n*−1^ for some *k*. If *k* ∈ **Z**_*i*_, then *k*–th coordinate of the sum 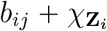 will be zero, thus reducing the number of 1 entries by 1. If *k* ∉ **Z**_*i*_, then the number of 1s in 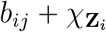 will increase by 1. □

Now we evaluate the sizes of sets in the matrix *M* using the sizes *κ*_*j*_ defined in Lemma 2.11.

##### Lemma 5.4.

*Consider* |**Z**_*i*_| = *m. Then, for any i* = 1, …, *n, the numbers in the i*–*th column of the matrix M are*

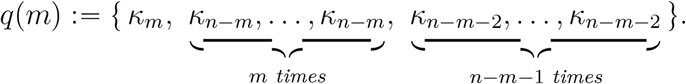

*Proof*. For vectors in B^*n*−1^ the possible sizes of sets *U* (*b*), *L*(*b*) are in the set {*κ*_0_, …, *κ*_*n*−1_}. The size of the set 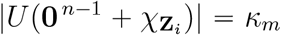. By Lemma 2.11, when 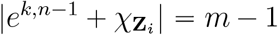 then the size of the set 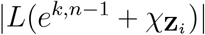 is *κ*_(*n*−1) − (*m*−1)_. By Lemma 5.3 this happens |**Z**_*i*_| = *m* times.

Similarly, By Lemma 2.11.2, when 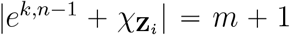 then the size of the set 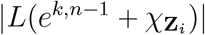 is *κ*_(*n*−1) −(*m*+1)_. By Lemma 5.3 this happens |**Z**_*i*_| = *n* − *m* − 1 times. □

##### Theorem 5.5.

*Consider network N* = (*G, δ*) *with n nodes, all-to-all coupling without self-edges and the sign-change function β* = (*β*_1_, …, *β*_*n*_) *induced by δ. Then the sign structure δ will produce an equipotent network if and only if the matrix M with numbers in column j induced by the choice of β*_*j*_ *is such that the product in each row is the same*.

In view of Theorem 5.5 to understand the network structures that support equipotency, we need to construct matrix *M* in (7) with equal products in all rows. This can be achieved by following these four steps:

##### Algorithm 5.6.

1. Determine the collections of numbers *K*_*n*−1_; these depend on the number *n* −1 of inputs to node *i*.
2. For every *i*, consider each potential size of the negative inputs

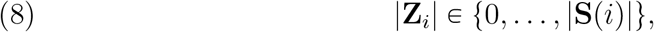

and compute the set of potential columns *q* (|**Z**_*i*_|) in matrix *M* .
3. Construct all matrices *M* from the columns *q* (|**Z**_*i*_|) in such a way that all rows have the same product. In the examples we compute below, this is achieved when |**Z**_*i*_|= |**Z**_*j*_| for all nodes *i, j* and when the product of elements in column *q* (|**Z**_*i*_ |)is maximal across all potential sizes in (8). This set of matrices corresponds to the collection of all maximal equipotent networks.
4. Among this set, find the set of non-isomorphic maximal equipotent networks.

This algorithm works for arbitrary networks. The main bottleneck for an arbitrary number of inputs *s* is computing the sets *K*_*s*_ that contain the sizes of the sets *U* (·) and *L* (·) in *MBF s*, as described in the remark after Lemma 2.11. In addition, if the network is not fully connected, then the set *K*_*s*_ for each node will depend on the network. When each column of matrix *M* has entries selected from a different set *K*_*s*_, equipotency would not be expected. Therefore, we restrict ourselves to networks *n* = 3, 4, 5 nodes with no self-edges where the number of inputs into each node is the same and, respectively, 2, 3, and 4. In these cases, we can compute the numbers in *K*_*s*_ explicitly,

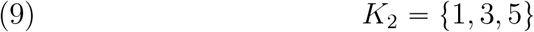

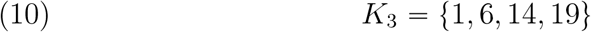

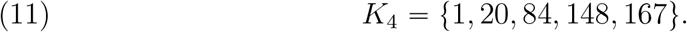

*Example: n =* 3. We follow the steps outlined in Algorithm 5.6. Using (9) and the fact that

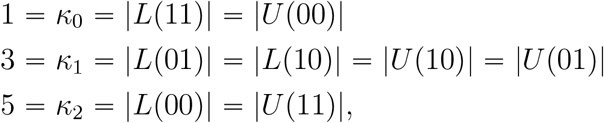

we evaluate columns in matrix (7). As result, we get three possible columns *q*(0), *q*(1), *q*(2) of matrix *M* listed in Table 1.

**Table 1.** The size of the set |**Z**_*i*_| on the entries in column of matrix *M* for different sizes of network *n*. The numbers in the third column are listed in order in lemma 5.4.

| $n$ | $\vert \mathbf{Z}_i\vert $ | $q(\vert \mathbf{Z}_i\vert )$ | |
| --- | --- | --- | --- |
| 3 | 0 | $\{\kappa_0, \kappa_1, \kappa_1\}$ | $\{1, 3, 3\}$ |
| | 1 | $\{\kappa_1, \kappa_2, \kappa_0\}$ | $\{3, 5, 1\}$ |
| | 2 | $\{\kappa_2, \kappa_1, \kappa_1\}$ | $\{5, 3, 3\}$ |
| 4 | 0 | $\{\kappa_0, \kappa_2, \kappa_2, \kappa_2\}$ | $\{1, 14, 14, 14\}$ |
| | 1 | $\{\kappa_1, \kappa_3, \kappa_1, \kappa_1\}$ | $\{6, 19, 6, 6\}$ |
| | 2 | $\{\kappa_2, \kappa_2, \kappa_2, \kappa_0\}$ | $\{14, 14, 14, 1\}$ |
| | 3 | $\{\kappa_3, \kappa_1, \kappa_1, \kappa_1\}$ | $\{19, 6, 6, 6\}$ |
| 5 | 0 | $\{\kappa_0, \kappa_3, \kappa_3, \kappa_3, \kappa_3\}$ | $\{1, 148, 148, 148, 148\}$ |
| | 1 | $\{\kappa_1, \kappa_4, \kappa_2, \kappa_2, \kappa_2\}$ | $\{20, 167, 84, 84, 84\}$ |
| | 2 | $\{\kappa_2, \kappa_3, \kappa_3, \kappa_1, \kappa_1\}$ | $\{84, 148, 148, 20, 20\}$ |
| | 3 | $\{\kappa_3, \kappa_2, \kappa_2, \kappa_2, \kappa_0\}$ | $\{148, 84, 84, 84, 1\}$ |
| | 4 | $\{\kappa_4, \kappa_1, \kappa_1, \kappa_1, \kappa_1\}$ | $\{167, 20, 20, 20, 20\}$ |

The highest product of the elements along the vector is 5 ∗ 3 ∗ 3 = 45 for column *q*(2); for *q*(0) the product is 9 and for *q*(1) the product is 15.

When all columns of matrix *M* are of type *q*(2) i.e. when |**Z**_*i*_| = 2 for *i* = 1, 2, 3, then

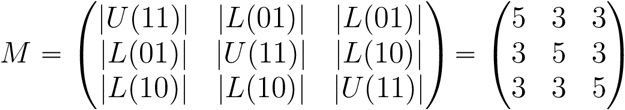

is an equipotent matrix. The corresponding equipotent network is in Figure 2a. The same observation applies to situation when |**Z**_*i*_ | = 0 for all i; the resulting matrix is

**Figure 2.**
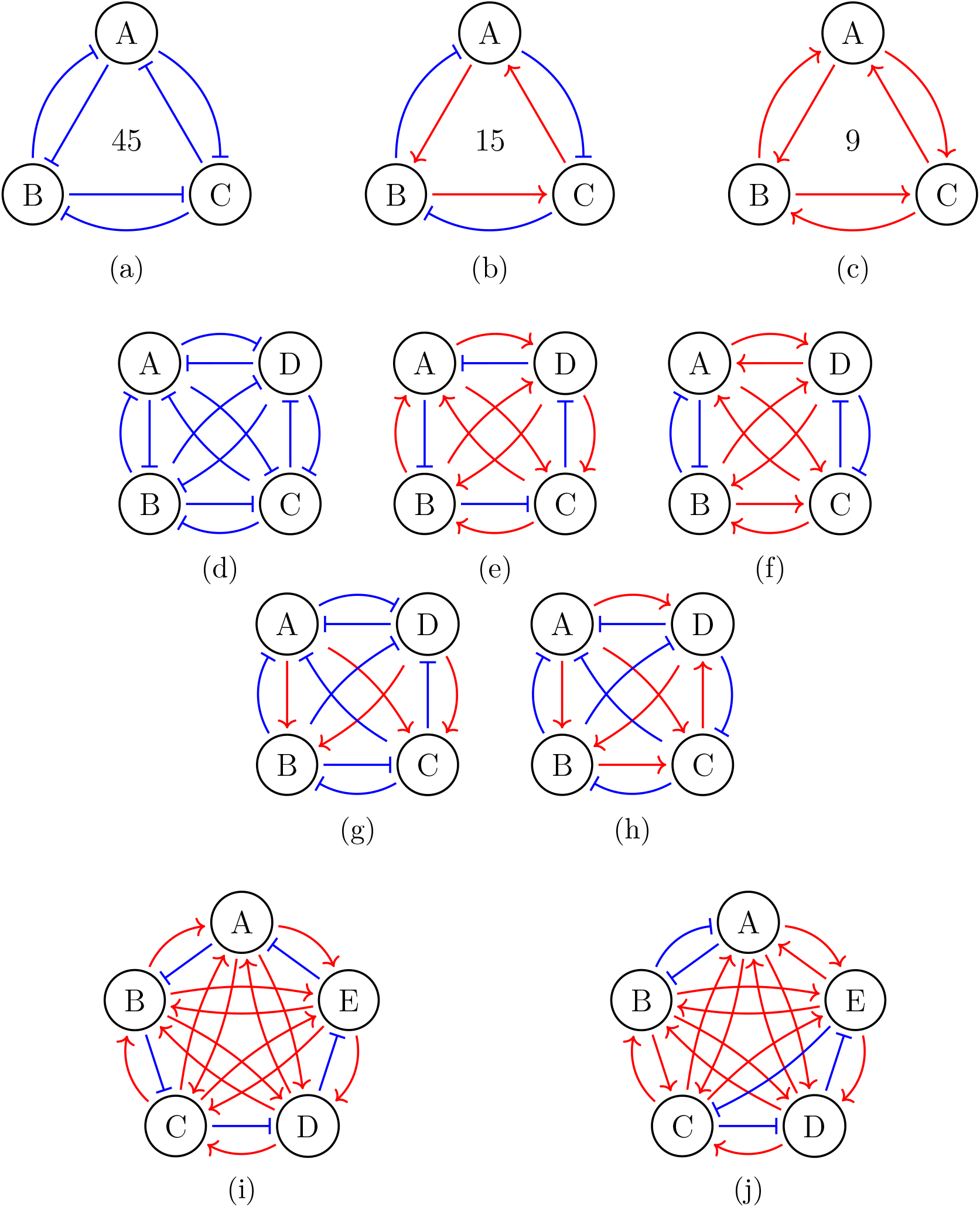
Network structures that support equal prevalence of single high states (equipotent networks). (a)-(c) equipotent three node networks. The number of MBMs supporting each single high state is indicated. (d)-(h) equipotent four node networks with maximum prevalence (4104 MBMs) among equipotent networks, (i)-(j) equipotent five node networks with maximum prevalence (1,979,631,360 MBMs) among equipotent networks.

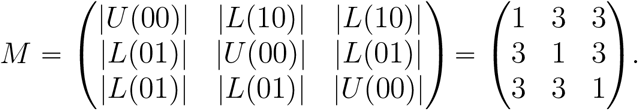

However, to anticipate the difficulties that are present for *n* ⩾ 4, not all choices of inputs when |**Z**_*i*_| = 1 produce an equipotent network. A non-symmetric network 2 ⊣ 1, 2 ⊣ 3, 3 ⊣ 2 produces matrix

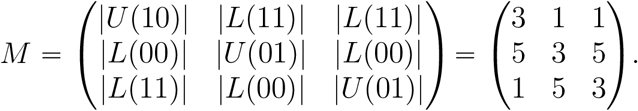

which is not equipotent. On the other hand, the symmetric network in Figure 2b is equipotent with matrix

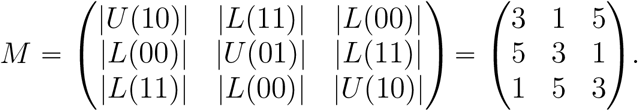

Finally, we show that if |**Z**_*i*_| ≠ |**Z**_*j*_|for some *i j*, then the network cannot be equipotent.

Consider the set *Z* : = {**Z**_1_, **Z**_2_, **Z**_3_} and let *η*_*i*_ : = |{*j* | |**Z**_*j*_| *= i* }| be the number of times that set of size *i* have been chosen. Then clearly *η*_0_ + *η*_1_ + *η*_2_ = 3. Counting how many times the column *q* (*i*), *i =* 0, 1, 2 contains elements 1 and 5 we note that number of 1s in matrix *M* is *η*_0_ + *η*_1_ and total number of 5s is *η*_1_ + *η*_2_. If the matrix is equipotent, then the number of 1s in each row must be the same, and the number of 5s in each row must be the same.

Since there are 3 rows in matrix *M*, both numbers *η*_0_ + *η*_1_ and *η*_1_ + *η*_2_ must be divisible by 3. Since 0 ⩽ *η*_0_ + *η*_1_ ⩽ 3 and 0 ⩽ *η*_1_ + *η*_2_ ⩽ 3, each sum must be either 0 or 3. Finally, because *η*_0_ + *η*_1_ + *η*_2_ = 3, there are only three possibilities

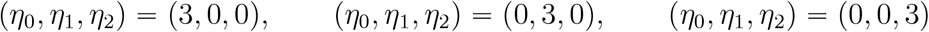

Thus, in an equipotent network, we must have |**Z**_1_| = |**Z**_2_| = |**Z**_3_| .

We summarize results in this section in a Theorem.

##### Theorem 5.7.

*Consider a* 3*-node network N with all-to-all coupling and no self-edges. Then N is equipotent if, and only if, for all i =* 1, 2, 3, *one of the following holds:*

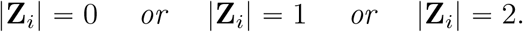

*Moreover,in the first case there are* 9, *second case* 15, *and third case* 45 *models supporting each equilibrium e*^1^, *e*^2^, *e*^3^.

Three equipotent networks are represented in Figures 2a to 2c, respectively.

*Example: n =* 4. We consider a network with 4 nodes and all-to-all coupling with no self-edges. We order the nodes 1, 2, 3, 4. Since each node receives input from the other three nodes, the set of inputs for each node *i* is **S**(*i*) = {1, 2, 3}.

We again follow the outline in Algorithm 5.6. Using (10) and the fact that

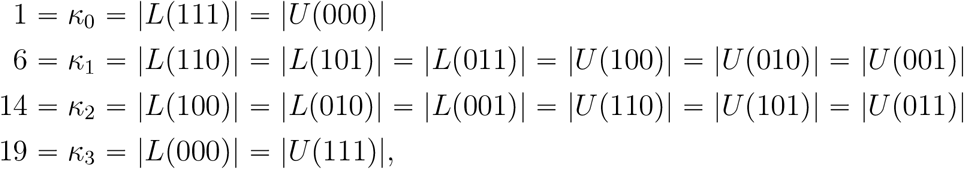

we evaluate columns in matrix (7). The four possible columns *q* (0), *q* (1), *q* (2), *q* (3) listed in Table 1.

We now encounter a degeneracy for *n* = 4, since there are only two (rather than four) distinct vectors *q*(·):

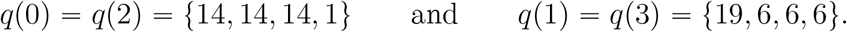

Let *C*_1_ = {0, 2} and *C*_2_ = {1, 3} be two classes of sizes of the input set |**Z**_*i*_| that produce two different vectors *q*.

As for the case *n =* 3, one can show that the only way to construct the equipotent matrix *M* is to select all four columns from the same class.

##### Corollary 5.8.

*Consider a* 4*-node network N with all-to-all coupling and no self-edges. Then N is equipotent if, and only if, either*

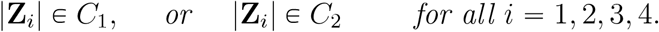

*In this first case (Type 1), there are* 2744 = 1 ∗ 14^3^ *and in the second case (Type 2)*, 4104 = 19 ∗ 6^3^ *models supporting each equilibrium e*^1^, …, *e*^4^.

We now construct all equipotent matrices *M* that yield the equipotent network with value 4104. Note that here, unlike the case *n =* 3, there are several different equipotent matrices *M* and hence different equipotent networks. By Corollary 5.8 the sets **Z**_*i*_ must satisfy |**Z**_*i*_| ∈ {1, 3} which means

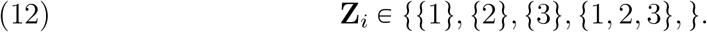

These choices are not independent. Observe that the choice of **Z**_*i*_ implies that in column *i* of matrix *M* there is a unique row index *r* (*i*) such that the entry *m*_*r (i)i*_ *=* 19, while the other entries are all 6. It is easy to check that each of the four choices above produces a different value of *r* (*i*) ∈ {1, 2, 3, 4} . In order to achieve equipotency, we must have that

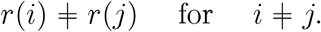

To construct such matrix *M* we proceed as follows

- For *i =* 1 we can select **Z**_1_ to be any of the 4 choices in (12);
- **Z**_2_ is then selected in such a way that *r* ≠ 2 *r* (1), which yields 3 choices. item **Z**_3_ is then selected in such a way that *r* (3) ≠ *r* (1), *r* (3) ≠ *r* (2) which yields 2 choices.
- **Z**_4_ is then uniquely determined by requirement that *r*(4) ≠ *r*(1), *r*(2), *r*(3).

Therefore, the number of equipotent matrices (networks) with value 4104 is at most 4 3 2 1 4! 24.

The same permutation argument applies to {14, 14, 14, 1}, which corresponds to |**Z**_*j*_| ∈ {0, 2} . This case also yields at most 4! 24 possible equipotent networks.

In the final step, we determine non-isomorphic equipotent networks within the collection of 24 networks, since some of the 24 equipotent networks may be equivalent under relabeling of the nodes. This is implemented in Python by generating all permutations of the node labels and selecting the lexicographically smallest relabeled tuple as the canonical form. Networks that have the same canonical form are placed in the same isomorphism class. In each equipotent class that contains 24 network we find 5 non-isomorphic networks, which are shown in Figures 2d to 2h.

*Example: n =* 5. Extending the previous construction for *n =* 4, we consider five nodes 1, 2, 3, 4, 5 with all-to-all coupling and no self-edges. Each node receives input from the other four nodes and thus for each node *i* the set of inputs is **S (***i*) *=* {1, 2, 3, 4}.

We again follow the outline in Algorithm 5.6. Using (11) and

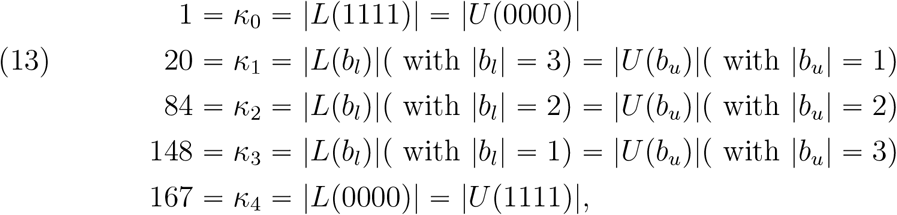

the five possible sizes of the set of negative inputs |**Z**_*i*_| ∈ {0, 1, 2, 3, 4} produce the five distinct columns *q* (|**Z**_*i*_|) of matrix *M* listed in Table 1. Thus, unlike the *n* 4 case, where two distinct types of *q* (|**Z**_*i*_|) occur, here every 5 value of |**Z**_*i*_| produces a distinct vector *q* (|**Z**_*i*_|) . As before, one can verify that any equipotent matrix *M* must have all columns from the same class. i.e., it is necessary that |**Z**_*i*_| *=* |**Z**_*j*_| = *m* for all *i, j* ∈ {1, …, 5}, and *m* ∈ {0, 1, 2, 3, 4}.

From Table 1 among these five possible values of *m*, the maximal product occurs when |**Z**_*i*_| = 1 and the product is

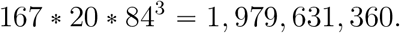

To construct the collection of equipotent matrices *M* where every column is *q* (1), we first observe that the diagonal entry in matrix (7) is 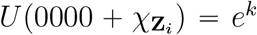 for some *k*. It follows from (13) that |*U* (*e*^*k*^)| = 20 and all the diagonal entries of matrix *M* are all equal to 20. Straightforward combinatorial calculation shows that there are 44 equipotent matrices *M* where each column is of type *q* (1) .

As in the case *n =* 4, some of these networks are the same after relabeling of the nodes. Grouping these 44 networks according to their canonical forms under relabeling of the nodes, produces 2 non-isomorphic equipotent networks, pictured in Figures 2i and 2j.

### 5.2 Networks with all-to-all coupling and fixed-sign of self-edges

In this section, we link the analysis of equipotency in networks with all-to-all coupling without self-edges to equipotency in networks with all-to-all coupling and fixed-sign of self-edges (either all self-activations or all self-inhibitions).

The main observation is that in the network with *n* nodes and all-to-all coupling with self-activation on each node, the set of negative inputs to node *i* is **Z**_*i*_ ⊂ **S**(*i*)\{*i*}. Because the self-edge is activating, the diagonal entries *U* p*b*_*u*_q in the matrix *M* in (7) will have *b*_*u*_ ≠ **0** but will have at least one entry (corresponding to the self-input) equal to 1 i.e. |*b*_*u*_| ⩾ 1 and |*U* (*b*_*u*_)| ∈ {*κ*_1_, *κ*_2_, …, *κ*_*n*_} ⊂ *K*_*n*_. By the same argument, the off-diagonal entries *L*(*b*_*l*_) will have *b*_*l*_ ≠ **1** and therefore the size 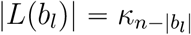 will also belong to the set {*κ*_1_, *κ*_2_, …, *κ*_*n*_} ⊂ *K*_*n*_.

It follows that for set of sizes 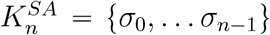 for network with all-to-all coupling with self-activations is related to set of sizes *K*_*n*_ = {*κ*_0_, …, *κ*_*n*_} by

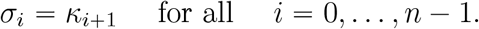

This implies that

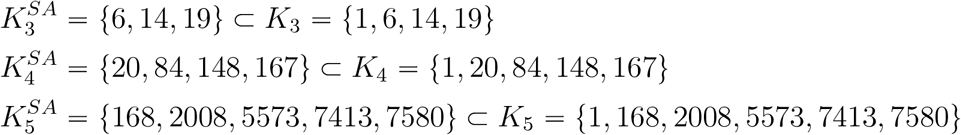

By a similar argument for a network with *n* nodes and all-to-all coupling with self-inhibition on each node, we will have *i* ∈ **Z**_*i*_. In this case, the diagonal entries *U* (*b*_*u*_) will have |*b*_*u*_| ⩽ *n* − 1 while the off-diagonal entries *L*(*b*_*l*_) will have |*b*_*l*_| ⩾ 1. Therefore the size of |*U* (*b*_*u*_)| and |*L*(*b*_*l*_)| ∈ {*κ*_0_, *κ*_1_, …, *κ*_*n*−1_} ⊂ *K*_*n*_. Therefore the set of sizes 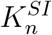 for networks with all-to-all coupling and self-inhibitions is

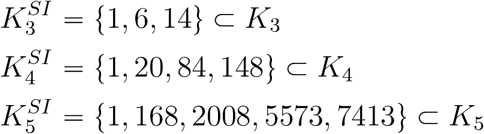

With these modifications, the Algorithm 5.6 can now be followed. The equipotent networks with self-edges would have the same sign-structure for the cross-couplings as networks without self-edges. Surprisingly, we find that the highest equipotency also occurs for the same number of negative inputs to each node, two for *n =* 3, one or three for *n =* 4, and one for *n =* 5. Therefore, the networks in Figure 2 (with added self-edges) still have the highest equipotency in their category. However, since 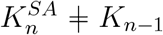 and 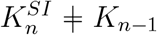 we do not know if this is true for arbitrary *n*.

## 6. Multistability of single high states

In this section, we ask a different question than in the previous section. Consider a collection of states *e*^1^, …, *e*^*n*^ ⊂ B^*n*^. While in the previous section we asked which networks have the property that the number of functions supporting *e*^*i*^ is the same as the number of functions supporting *e*^*j*^, for all *i, j*, here we focus on the existence of a single MBM that is capable of supporting *all e*^1^, …, *e*^*n*^ ⊂ B^*n*^.

Our main result in this section is the following Theorem.

### Theorem 6.1.

*Consider a network N* = (*V, E*) *with* |*V*| = *n nodes. Then N supports full multistability if and only if each vertex i either has a positive self-edge, or all other vertices have repressive edges terminating at i*.

Note that the networks that support full multistability include the *n*-team network, where each vertex forms its own team. Such a network has positive self-edges on each node and all other edges are repressive and thus satisfy both conditions of the Theorem at the same time. However, the Theorem shows that one of these conditions is sufficient for full multistability: if positive a self-loop exists, the character of incoming edges is irrelevant. A natural conjecture is whether, in this case, <u>the number</u> of MBFs supporting full multistability is affected by the character of incoming edges and whether the n-team network has the maximal number of MBFs supporting full multistability among all such networks. As we show later, this is indeed true for *n* 3, 4, but surprisingly not true for *n* 5. The general problem of network structure that supports full multistability for the maximal number of compatible functions remains open.

In order to prove 6.1, we first prove several Lemmas.

### 6.1. Networks without self-edges

#### Lemma 6.2.

*Consider network N* = (*V, E*). *Assume that for all i* “1, …, *n, i* ∈ **S**(*i*), *i.e*., *that the network N does not have a self-edge*.

*If N supports full multistability then for all i* = 1, …, *n*,

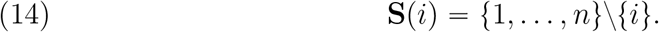

*In other words, the network has to have all-to-all coupling*.

*Proof*. Let *δ* be the sign function and *β* = (*β*_1_, …, *β*_*n*_) the corresponding change of sign function. Without loss of generality, assume *i* = 1, and that node 1 has inputs from the next *s* nodes with *s +* 1 < *n*. The evaluation of sources {2, …, *s +* 1} on steady states (*e*^1^, *e*^2^, …, *e*^*s*+1^, …, *e*^*n*^) produces the following set of input vectors in B^*s*^

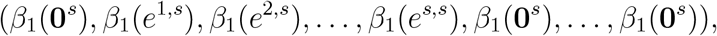

where **0**^*s*^ is the zero vector and *e*^*j,s*^ is the unit vector in *j–*th direction in B^*s*^. The terms **0**^*s*^ at the end correspond to evaluation of sources on steady states *e*_*j*_ with *j* = *s +* 1, …, *n*;

By Theorem 3.5 if *f* = (*f*_1_, …, *f*_*n*_) supports multistability, then

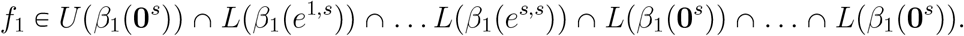

Note that this intersection contains the terms

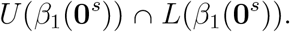

Since *U* (*b*) ∩ *L* (*b*) = ∅ for any *b*, the intersection is empty. Since *β* was arbitrary, if *v*_1_ has *k +* 1 < *n* inputs, no sign structure *δ* can support full multistability between *e*^1^, …, *e*^*n*^. Since *v*_1_ was arbitrary, if *N* supports full multistability then (14) holds. □

Therefore, networks that can possibly support full multistability without self-edges must have all-to-all coupling.

In order to understand what types of sign structure on *N* with all-to-all coupling support full multistability, we will investigate in more detail the effect of the sign function *β*_*i*_ associated with a vertex *i* with sources *j* P **S**(*i*) and |**S**(*i*)| = *n* − 1.

#### Lemma 6.3.

*Consider network N* = (*V, E*) *where each vertex i receives input from all other nodes, except itself. Consider vertex i and consider an arbitrary change of sign function β*^**Z**^ *at vertex i for some set of negative inputs* **Z** ⊂ **S** (*i*).

*Then all MB functions f*_*i*_ *at node i that support full multistability satisfy*

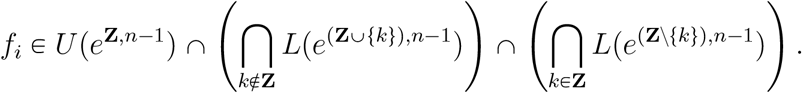

*Proof*. Consider all steady states *e*^*k*^, *k* = 1, …, *n* which must be supported by the function *f*_*i*_ at node *i*. We now describe conditions that arise from each steady state *e*^*k*^. We discuss three cases:

1. *k* = *i*;
2. *k* ≠ *i, k* ∉ **Z**.
3. *k* ≠ *i, k* ∈ **Z**;

In the first case, the *i*–th coordinate of *e*^*k*^ is one, and so *f*_*i*_ ∈ *U* (·) and 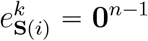.

As defined, 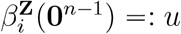 where

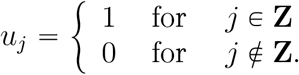

It follows that the first case leads to the condition *f*_*i*_ *i*–th *U* (*e*^**Z**^).

In the second and third cases, the *i*–th coordinate of *e*^*k*^ is zero, and so *f*_*i*_ *i*–th *L*(·) and 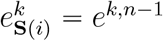. However, the input into the function *L* is different between the second and third cases.

For the second case, since *k* ∉ **Z**, the value remains the same: 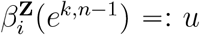 with

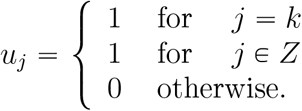

We set *β*_*i*_ (*e*^*k,n*−1^ = *e*^**Z′**,*n*−1^, **Z′ = Z** ∪ {*k*} . If |**Z**| *m*, then there are *n* − *m* − 1 steady states *e*^*k*^ in this category.

In the third case, since *k* ∈ **Z**, the value flips: 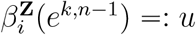

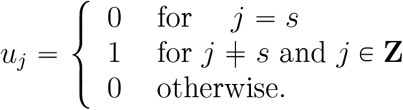

Set *β*_*i*_ (*e*^*k,n*−1^ = *e*^**Z′**,*n*−1^, **Z′ = Z**\{*k*} . There are *m* steady states *e*^*k*^ in this category. Cases 1,2 and 3 correspond to three terms in the statement of the Lemma. This concludes the proof. □

#### Theorem 6.4.

*Consider network N* = (*V, E*) *where each vertex i receives input from all other nodes, except itself*.

*If N supports full multistability, then for all i* = 1, …, *n, every component* 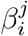 *of the the function β*_*i*_ *is* 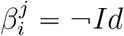.

*Proof*. By Lemma 6.3

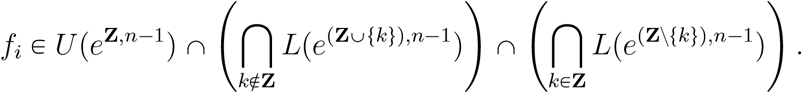

Assume **Z ⊊ S** (*i*) and recall that |**S**(*i*) = *n* − 1. Then there is *j* ∉ **Z**, and therefore there is at least one term in the intersection of the form *L*(*e*^(**Z**∪{*j*}),−1^). Consider now the sets *U*(*e*^**Z**,*n*^) and *L*(*e*^(**Z**∪{*k*}),−1^). Since *e*^**Z**,*n* −1^ ≺ *e*^(**Z**∪{*j*}),−1^ and *f*_*i*_ are increasing monotone Boolean function we must have

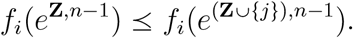

However, the requirement that *f*_*i*_ ∈ *U* (*e*^**Z**,*n* −1^) ∩ *L* (*e*^(**Z**∪{*j*}),−1^) implies that

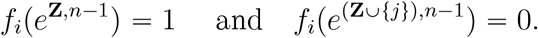

i.e, *f*_*i*_ ∈(*e*^**Z**,*n* −1^) ≻ *f*_*i*_(*e*^(**Z**∪{*j*}),−1^). This contradiction implies **Z = S**(*i*). □

We observe that if **S**(*i*) = *V\*{*i*} then the set of MBF *f*_*i*_ that supports full multistability satisfies

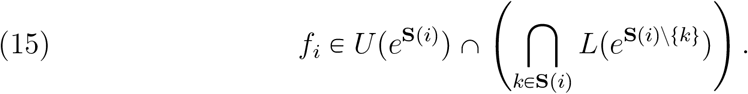

Figure 3a shows the only input structure to node *A* supporting full multistability in a 3-node network without self-loop on *A*.

**Figure 3.**
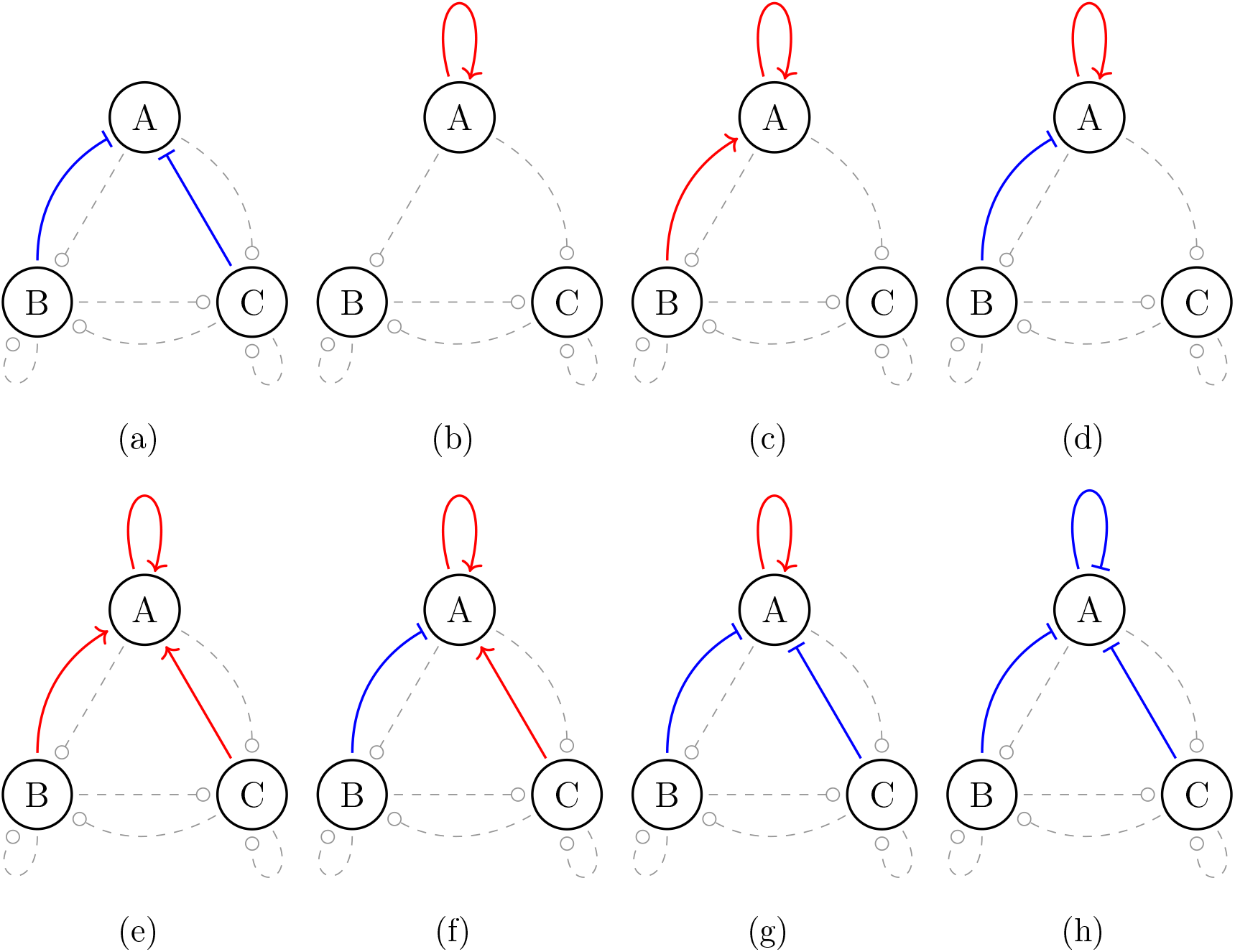
Network structures that support multistability of single high states. Input structures to node *A* in a 3-node network compatible with multistability between (100), (010), (001) . Dashed lines indicate that the input structures for nodes *B* and *C* could take any choice from the corresponding structures. (Isomorphic structures are not included)

#### Lemma 6.5.

*Consider network N* = (*V, E*) *where each vertex i receives input from all other nodes, except itself*.

*Then, for any node i, all MB functions f*_*i*_ *that support full multistability must be an essential monotone Boolean function*.

*Proof*. If *f*_*i*_ is non-essential MBF, there is an input *j* such that *f*_*i*_ is redundant in the input *j*

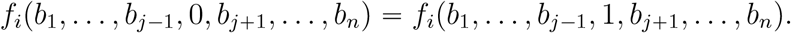

On the other hand, it follows from (15) that

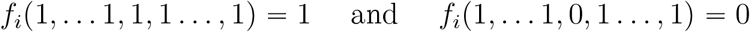

for any position *j* of the input 0. This contradiction shows that *f*_*i*_ is an essential monotone Boolean function. □

### 6.2. Networks with self-edges

We now consider networks with self-edges. Recall from Theorem 3.5 that *f* =(*f*_1_, …, *f*_*n*_) supports full multistability if and only if

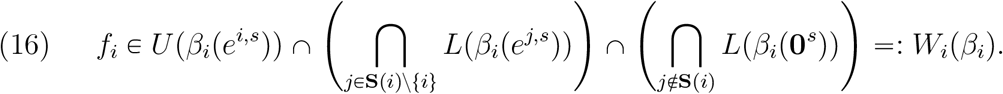

We fix vertex *i* and investigate the effect that the change of sign function *β*_*i*_ has on the set *W*_*i*_(*β*_*i*_).

#### Lemma 6.6.

*If β*_*i*_ *satisfies*

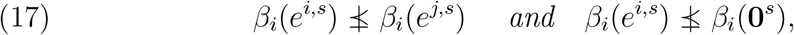

*then*

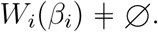

*If at least one of the conditions in* (*17*) *is not valid, i.e. either there exists j such that β*(*e*^*i,s*^) ≺ *β*(*e*^*j,s*^) *or β*(*e*^*i,s*^) ≺ *β*(**0**^*s*^), *then*

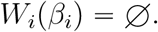

*Proof*. Since *f*_*i*_ is an increasing MBF, the fact that *f*_*i*_ ∈ *U* (*β*_*i*_(*e*^*i,s*^)) implies that

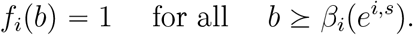

On the other hand the requirement that *f*_*i*_ ∈ *L* (*β*_*i*_(*e*^*i,s*^)) implies that

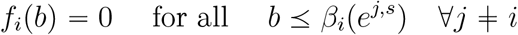

and *f*_*i*_ ∈ (*L*(*β*_*i*_(**0**^*s*^)) implies that

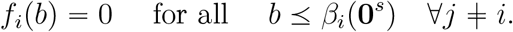

Define the set

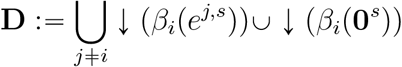

and **U** := ↑*β*_*i*_(*e*^*i,s*^). Then the conditions (17)

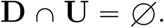

Therefore, any function *f*_*i*_ whose truth set T ⊃ **U** includes **U** and whose kernel *ker f* ⊃ **D** includes **D**, supports full multistability.

To prove the second statement, we note that if there exists *j* such that *β*_*i*_(*e*^*i,s*^) ≺ *β*_*i*_(*e*^*j,s*^) then

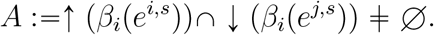

Then for *b* ∈ *A* any *f*_*i*_ ∈ *W*_*i*_(*β*_*i*_) must satisfy *f*_*i*_(*b*) = 1 and also *f*_*i*_(*b*) = 0. Therefore *W*_*i*_(*β*_*i*_) = ∅. Similar argument applies to the statement *β*_*i*_(*e*^*i,s*^) ≺ *β*_*i*_(**0**^*s*^). □

If *a, b* ∈ B^*s*^ are not comparable, we say

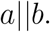

**Example**. If *β*_*i*_ = (*Id*, …, *Id*) and thus all edges are positive then the conditions of Lemma 6.6 read

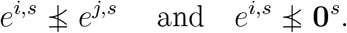

Since *e*^*i,s*^||*e*^*j,s*^ for all *j* ≠ *i* and *e*^*i,s*^ ≻ **0**^*s*^ these conditions are satisfied.

#### Lemma 6.7.

*Consider any change of sign function* 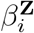 *on* B^*s*^. *If i, j* ∉ **Z**, *then* 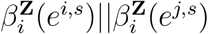 *for all j* ∉ *i and* 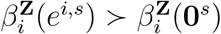.

*Proof*. Since *e*^*i,s*^ || *e*^*j,s*^ and *i*, ∉*j* **Z** then *i* and *j* components still remain non-comparable after applying 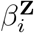. Since *e*^*i,s*^ ≻**0**^*s*^, it is easy to see that adding a series of 1s to both vectors does not change their order. □

#### Lemma 6.8.

*Consider node i with s inputs, including from i. If the self-loop i* ⟶ *i is positive, then for any change of sign function* 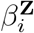 *on* B^*s*^ *the set* 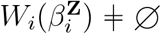 .

*Proof*. Our assumption implies that *i* ∉ **Z**. If *j* ∉ **Z** then by previous Lemma 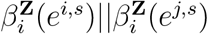 for all *j* ≠ *i*.

Consider *j* ∈ **Z**. Note that *β*_*i*_(*e*^*i,s*^) ≺ *β*_*i*_(*e*^*j,s*^) since the *j*–th component of *β*_*i*_(*e*^*j,s*^) is zero and *j*–th component of *β*_*i*_ (*e*^*i,s*^) is one, and on all other components the vectors agree.

Therefore, using the previous Lemma, all assumptions of Lemma 6.6 are satisfied and 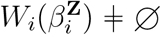. □

Figures 3b to 3g shows the possible input structures to node *A* supporting full multistability in a 3-node network with self-activation on *A*. Every structure, up to isomorphism, is listed. After appropriate relabeling of nodes, input structures to *B* and *C* can be chosen arbitrarily from this set. In the next Lemma, we denote by [*m*] = {1, …, *m*} the set of positive integers that are smaller than or equal to an integer *m*.

#### Lemma 6.9.

*Consider node i with s inputs, including from i. If the self-loop i* – *i is negative, then the set* 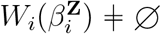 *if and only if s* = *n and* **Z** = [*n*].

*Proof*. If the self-loop *i* ⟶ *i* is negative, then *i* ∈ **Z**. If **Z** ⊊ [*s*] there is *j* ≠ *i* and *j* ∉ **Z**. Then 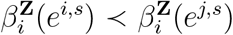 since 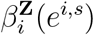 has component *i* equal to zero, while all components in **Z**\{*i*} are equal to 1. On the other hand, 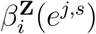 has all components in **Z** equal to 1, including the component *i*. By Lemma 6.6 the set 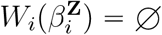.

If **Z** = [*s*] but *s* ≠ *n* then vector 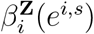 has all entries 1 except entry which is 0. On the other hand, 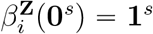. Therefore 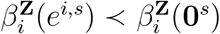 and by Lemma 6.6 the set 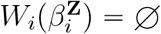.

Finally, if *s* = *n* and **Z** = [*n*], then 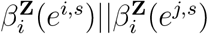 are not comparable. The term 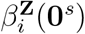 is not present in equation (16). Then by Lemma 6.6 the set 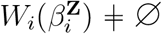.

As a consequence of this Lemma, there is only one input structure that supports full multistability at a node with self-inhibition. An example in a 3-node network is depicted in Figure 3h.

#### Remark 6.10.

If a vertex *i* has a positive self-loop, the set of functions supporting full-multistability includes the function *f*_*i*_ ≡ *b*_*i*_, which does not depend on any other input, and therefore the inputs from all other vertices are redundant. So this class of functions contains the class of completely disconnected networks discussed in Corollary 4.4

On the other hand, if a vertex *i* has a negative self-loop, then MBF that supports multistability, the input from *i* must be redundant.

#### Theorem 6.11.

*Consider a fully-connected network with self-edges N* = (*V, E*) . *If i had a negative self-loop, then the only monotone Boolean function that supports full multistability is*

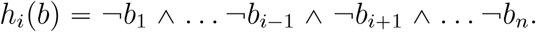

*Proof*. Fix *i* and assume *i* had a negative self-loop. In view of Lemma 6.9, if *h*_*i*_ supports full multistability, then *h*_*i*_ has negative inputs from all nodes **Z = [***n*] . Recall that 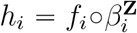 and *f*_*i*_ is positive monotone Boolean function *f*_*i*_ ∈ *MBF*(*n*) . We denote by 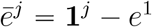 a vector of ones with a single zero at position *j*.

Then the set functions *f*_*i*_ that support *e*^1^, …, *e*^*n*^ lie in the intersection

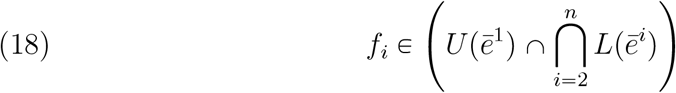

We show that the intersection in this case only contains a single MB function *f* with truth set

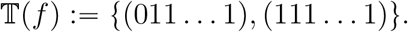

Let *Q* := {(011 … 1), (111 … 1)}. The fact that *Q* ⊂ T(*f*) follows from the requirement that *f*_*i*_ ∈ *U* (011 … 1) and monotonicity of *f*_*i*_.

To show the other inclusion, consider the value of *f*_*i*_ on *b* ∈ B^*n*^ stratified by the number of ones |*b*| in *b* ∈ B^*n*^.

- |*b*| = *n*. Then *b* “**1** and *f*_*i*_p*b*q = 1 by monotonicity of *f*_*i*_.
- |*b*| = *n* − 1. Then by (18), 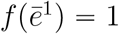 and 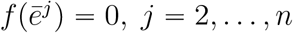
- |*b*| < *n* − 1. For any such *b* there is a vector *b*^′^ with |*b*^′^| = *n* − 1 and

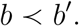

Since *f*_*i*_ P *L*(*b*^′^) it follows that *f*_*i*_(*b*^1^) = 0 by monotonicity of *f*_*i*_.

It follows that *f*_*i*_ (*b*) = 0 for all *b* ∉ *Q* and therefore from T *f* ⊂ *Q*.

The only function *f*_*i*_ with truth set *Q* is the function

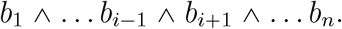

Theorem follows from **Z** “r*n*s. □

### 6.3. Networks maximizing prevalence of full multistability

An interesting question remains. When a node *i* has a positive self-edge and inputs from all other edges, what size of the set of negative inputs |**Z**_*i*_|to node *i* maximizes the number of MBFs that support multistability? On the first glance, one would conjecture that this happens either with |**Z**_*i*_| = 0 (i.e., all inputs are positive) or |**Z**_*i*_| = *n* − 1 i.e., all inputs from other nodes except the self are negative. However, the answer seems to be more subtle and depends on the internal structure of the lattices *MBF* (*s*) . We provide a few numerical examples that illustrate a somewhat surprising answer.

In particular, we use Algorithm 6.12 described below to count the number of functions that support multistability for fully connected networks with self-activations for a fixed value of *n* across varying numbers of inhibitory inputs |**Z**_***i***_| . Table 2 gives the calculations for *n* = 3, 4, 5 nodes using this algorithm with *W*_*i*_ being the number of monotone Boolean functions at node *i* supporting multistability. We can see that for *n* = 3, 4 the network with inhibition (|**Z**_***i***_| = *n* − 1 for every node) from every other node besides itself has the most monotone Boolean models supporting multistability. However, for *n* = 5, the network with |**Z**_*i*_| = 1 negative input for each node has the most models supporting multistability. At this moment, we lack any combinatorial theory that would explain this pattern and extend it to *n* > 5.

**Table 2.** Number of multistable functions for each node in a *n*-node fully-connected networks with fixed self-activations and |**Z**_***i***_| - inhibitions from other nodes.

| $n$ | 3 | | | 4 | | | | 5 | | | | |
| --- | --- | --- | --- | --- | --- | --- | --- | --- | --- | --- | --- | --- |
| $\vert \mathbf{Z}_i\vert $ | 0 | 1 | 2 | 0 | 1 | 2 | 3 | 0 | 1 | 2 | 3 | 4 |
| $\vert W_i\vert $ | 2 | 3 | 10 | 9 | 17 | 14 | 38 | 114 | 575 | 445 | 148 | 334 |

#### Algorithm 6.12.

Consider node *i*. The algorithm counts the number of monotone Boolean functions *f*_*i*_ with *n* inputs and *m* ⩽ *n* − 1 inhibitory inputs. We assume without loss of generality that the first *m* inputs excluding self-edges (**S** (*i*) *\* {*i*}) are inhibitory. Our main tool is the constraint (16) that determines which functions *f*_*i*_ support full multistability.

We follow the following steps:

1. We find the value *b*_*u*_ such that *U* (*b*_*u*_) is a component of the intersection in (16) and collect all values *B* := {*b*_*l*_} such that *L*(*b*_*l*_) is a component of the intersection in (16). The function *f*_*i*_ must satisfy

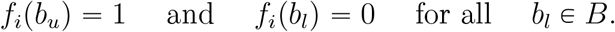
2. We use the constraint that *f*_*i*_ must be a monotone function. Therefore,

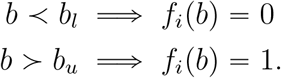
3. Let *C* ⊂ B^*n*^ be the set of vectors that have been assigned a value in steps (1) and (2). Consider the partially ordered set *P* := B^*n*^\*C*. This is the set of *unconstrained* vectors. Assume *p* := |*P* | be the size of *P* .

Figures 4a to 4c illustrates the imposition of the constraints for *n* = 3 and number of negative inputs *m* = 0, 1, 2. Note that when *m* = 0, the set *P* = {(011)}, when *m* = 1 the set *P* = {(100) ≺ (101)} and when *m* = 2 the set *P* = {(100) ≺ {(100), (101), (011)}}.

**Figure 4.**
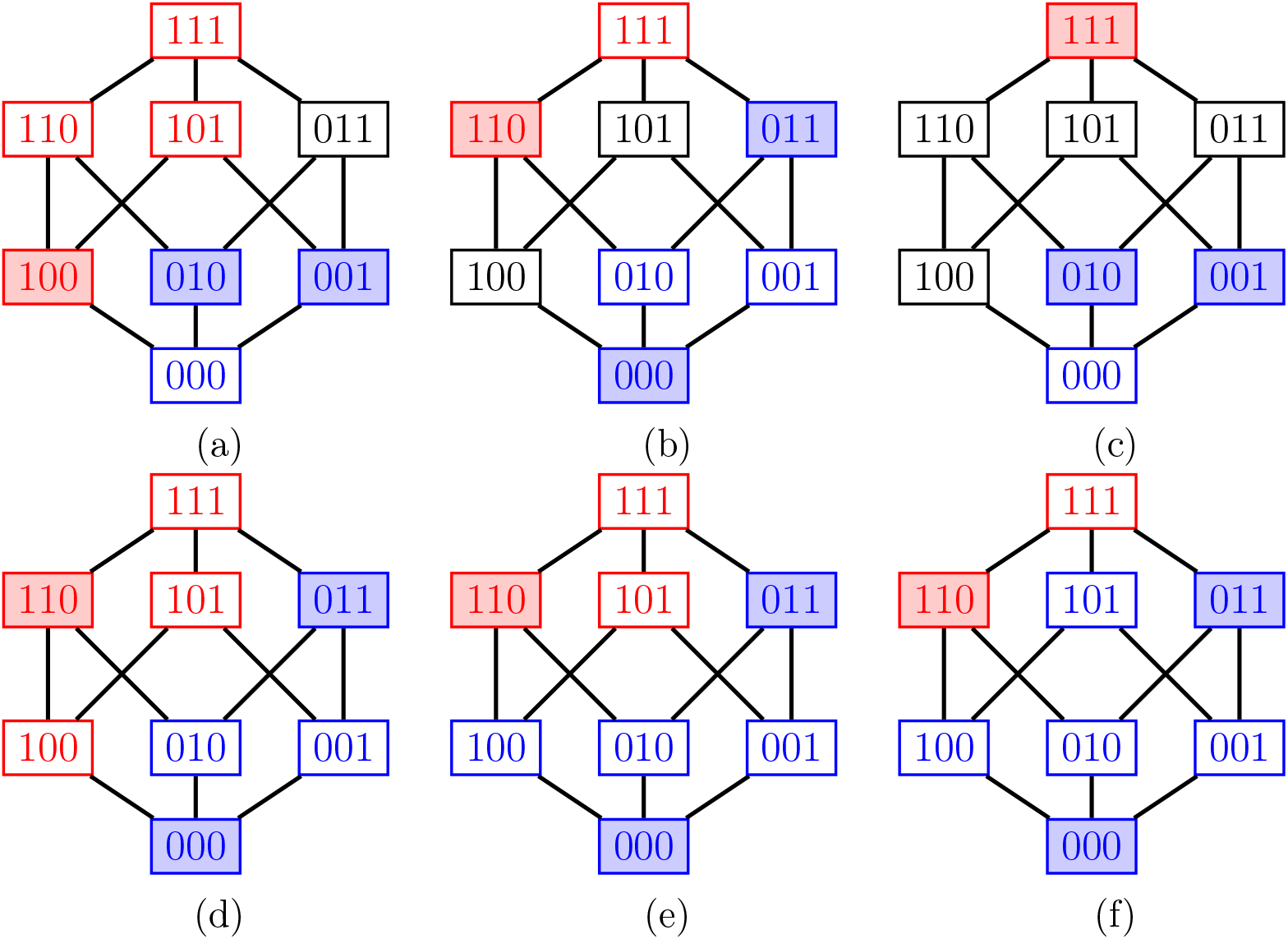
Counting functions *f*_*i*_ that support multistability between single-high states. Lattice of Boolean vectors *b ∈* B^3^ with constraints imposed by *U* (*β*(100)) ∩ *L*(*β*(010)) ∩*L*(*β*(001)) for |**Z**_*i*_| inhibitory inputs. (a): |**Z**_*i*_| = 0, (b): |**Z**_*i*_| = 1, (c): |**Z**_*i*_| = 2. Blue nodes indicates *b* for which *f*_*i*_ (*b*) = 0 and red indicated *b* for which *f*_*i*_ (*b*) = 1. The shaded nodes denote direct constraints, while values of *f*_*i*_ at nodes with a color outline are constrained by monotonicity. (d)-(f): Three possible configurations of *f*_*i*_ for |**Z**_***i***_| = 1 that follow monotonicity.

Given a set *P*, we proceed to recursively count monotone functions *f*_*i*_ that have values constrained on vectors *C*. Note that this number is bounded above by 2^*p*^, which would be achieved if vectors in *P* were unordered. When there are values *b, c P* such that *b* ≺ *c* are ordered, then monotonicity of *f*_*i*_ implies *f*_*i*_ (*b*) *⩽ f*_*i*_ (*c*) . This restricts the number of possible functions *f*_*i*_. Figures 4d to 4f show three functions *f*_*i*_ that are compatible by constrains imposed by set *P* = {(100) ≺ (101)} for *n* = 3 and *m* = 1.

1. Find the current poset *P* .
2. Select the set of highest nodes *H* in poset *P* .
3. Generate all binary assignments of values {0, 1} to nodes in *H*.
4. For each assignment, compute the set of vectors *C*^1^ ⊂ *P* that are newly constrained by the assigned values to *H* using monotonicity. Compute a new unconstrained poset *P* and return to step 1.
5. Repeat steps 1-4 until the set *P* is empty. Return 1 in this case.
6. Add the number of 1s across all iterations.

## 7. Discussion

In this paper, we study a structural question in systems biology of regulatory networks: which network structure is optimal for supporting the full set of single high states as stable steady states. These states are observed in developmental networks where each cell fate is associated with a single activated master regulator. Clearly, the optimality must be defined and will inevitably depend on the choice of the network model. We chose to address this question in the context of monotone Boolean models that are compatible with the regulatory network structure. The fact that we consider the collection of all such models derives from our desire to emulate ideas from continuous dynamical systems, where key insights are provided by the bifurcation theory. Bifurcation theory describes when the dynamics of the system qualitatively change and thus is intimately linked with questions of robustness of the dynamics. The key property that supports bifurcation theory is the smooth dependence of the dynamics on parameters.

In contrast, Boolean models are usually studied in isolation. However, there is a natural measure of closeness between Boolean maps based on the Hamming distance between their ranges, which allows studying changes in dynamics between nearby maps. This paper is thus a continuation of our program studying dynamics of the collection of all monotone Boolean functions compatible with the signed network structure [2, 15, 16] (see also [1] for early efforts in this direction.)

In the previous paper [2], we have characterized a set of monotone Boolean models *f* = (*f*_1_, …, *f*_*n*_) where each *f*_*i*_ is an increasing monotone Boolean function that supports a state *b* ∉ B^*n*^ as a steady state. This corresponds to monotone Boolean models with an influence network that has only positive edges. In the same paper, we have generalized this result to *balanced networks* which are directed graphs with no negative cycles. It is known that monotone Boolean models of balanced networks are related by a change of variables to models whose influence network has only positive edges.

The major theoretical advance in this paper is the characterization of all monotone Boolean models that support a state *b* ∈ B^*n*^ as a steady state for <u>arbitrary</u> influence networks. Importantly, such networks may also support complex attractors. For any *b* ∈ B^*n*^ we characterize a set of models *V*_*b*_ : = {*h* | *h (b)*= *b}* . If we denote by *MBM (N)* the set of monotone Boolean models *h* : B^*n*^ → *B*^*n*^ with given influence network *N*, then the set

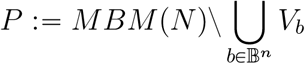

is non-empty and contains models *g* for which all attractors are complex. Direct characterization of *g* ∈ *P* remains an open problem.

We note that since the number of MBFs increases rapidly and is only explicitly known for *n* ⩽ 9, the scalability of numerical simulations is limited. However, our characterization provides a compact description of steady states and multistability for any network of any size *n*.

We first examine network structures that support all single high states with the highest possible number of monotone Boolean models. We show that the network that achieves this number is a network with isolated nodes with positive self-edges. Since such a network cannot represent a decision-making network, we study sign structures in networks with *n* = 3, 4, 5 nodes and all-to-all coupling that support maximal equipotency. Surprisingly, while for *n* = 3, 4 such networks have all negative incoming edges from other nodes, for *n* = 5 the networks that support maximal equipotency have exactly one negative in-edge, with all other edges positive. We note that several networks that support maximal equipotency are relatively far apart from each other in the space of networks, requiring a change of sign on several edges. So the landscape of degree of equipotency in the space of networks has several well-separated optima.

We then examine networks that support multistability among all single high states. Our main result shows that each node in such networks either has a positive self-regulation or all edges that terminate in it must be negative. Clearly, this class includes both networks with isolated nodes and positive self-edges, and fully connected networks where all mutual edges are negative. Since the first type of networks can be considered completely *structurally modular*, and the second type is most certainly not, this shows that the ability to support *functional modularity* of simultaneous stability of single high states does not depend on structural modularity.

It is important to contextualize these results within the complexity of real biological networks. Such networks often involve significant redundancy and additional layers of regulation by signaling molecules, such as morphogens or cytokines, that play a key role in the process of differentiation. In this study, however, we focus our analysis on the core of the network, specifically investigating the expected effective regulation among the master regulators. This represents a coarse-grained or “condensed” network architecture, which, while sacrificing some of the fine-grained dynamic behavior provided by intermediate interactions, considerably simplifies the analysis. As such, our study can be seen as analyzing densely connected motifs that may be part of larger hierarchical network structures [21]. How these smaller motifs are integrated and give rise to emergent properties observed in coarse-grained networks is a topic of ongoing research [3, 26].

A primary assumption in our analysis is that a single network architecture governs differentiation into *n* distinct cell fates. However, the process of differentiation to highly divergent lineages may involve context-specific interactions, especially among the intermediate regulations, altering the topology during intermediate stages of differentiation. By not explicitly incorporating irreversibility, which is characteristic of developmental systems, our results remain broadly applicable to systems with high plasticity, such as transdifferentiation and cell-state transitions in cancer. Beyond changes in network topology, the stability of these states can also be modulated by variations in regulatory strength. Epigenetic remodeling, for instance, can stabilize specific fates by modulating the effective strengths of regulatory edges [17]. While our formalism is limited to the presence or absence of activatory or inhibitory interactions, we can capture epigenetic remodeling by gradually excluding a specific subset of models (or by focusing on the complementary subset), effectively removing the influence of certain inputs [31].

It is important to connect our results to results that can be expected for differential equations (ODEs) models. On one hand, there is a direct correspondence between monotone Boolean models and the behavior of Glass type ODE models [8, 14, 18, 11, 10] with piece-wise constant nonlinearities. In fact, monotone Boolean model results are valid for an open subset of parameters for such ODEs [9] as well as their counterparts with sufficiently steep nonlinearities [12]. We therefore view the network structures identified by our framework as a conservative estimate of all network structures supporting a particular property.

However, connecting network structure with specific dynamical behavior within an ODE framework is hindered by two fundamental difficulties. One is the uncountability of the ODE parameter space, which makes it impossible to examine the set of compatible models; extensive parameter sampling can neither establish nor rule out the existence of particular dynamics. Despite this limitation, empirical explorations of network architectures from ODE simulations yield results that broadly align with the theoretical findings from our Boolean framework [23]. The second difficulty is the uncountability of the phase space, where identifying dynamics as “the same” for the purpose of their classification can only be approximate. Nevertheless, studies have conducted exhaustive explorations of the space of network structures, combined with clustering, to map properties of this space [28, 5, 24]. However, these methods are restricted to analysis of smaller motifs (*n* ⩽ 3 or 4) due to the computational complexity of the analysis and the fact that the set of potential network structures increases rapidly with *n*.

Our results are complementary to efforts to develop methodologies for identifying gene regulatory networks from data [22, 29]. Our theoretical understanding of network design principles may guide some of these efforts, which are challenged by high uncertainty and can both miss existing edges and produce spurious connections [20].

Our framework enables the design of networks with targeted functional properties in synthetic biology. While ODE-based modeling has shown significant promise in the design of synthetic circuits [25, 32], the scalability constrains its utility. A fundamental bottleneck for the broader implementation of our framework is its computational complexity. While the underlying mathematical theory of our framework is dimension-independent, current implementations are constrained by the difficulty of computing up-sets and down-sets in the lattice of monotone Boolean functions and the complexity of the Boolean satisfiability problem [7]. Further work is required to develop this tool to allow biologists to design complex circuits without the need to navigate the underlying mathematical formalism.

## Additional Information

### Author Contributions

HBV: Conceptualization, investigation, formal analysis, visualization, writing-original draft. SA: Investigation, formal analysis, visualization, writing-original draft. MKJ: Conceptualization, writing-reviewing and editing. TG: Conceptualization, formal analysis, writing-original draft, writing-reviewing and editing.

### Data Accessibility

The code used are available at: https://github.com/Harshavardhan-BV/Monotone-Bool-Net

### Declaration of AI use

Gemma4 was used to enhance clarity for parts of the Discussion. After using this tool, the authors reviewed and edited the content as needed and take full responsibility for the content of the publication.

